# Mapping the mechanical microenvironment in the ovary

**DOI:** 10.1101/2021.01.03.425098

**Authors:** Thomas I. R. Hopkins, Victoria L. Bemmer, Stephen Franks, Carina Dunlop, Kate Hardy, Iain E. Dunlop

## Abstract

Follicle development in the human ovary must be tightly regulated to ensure cyclical release of oocytes (ovulation), and disruption of this process is a common cause of infertility. Recent ex vivo studies suggest that follicle growth may be mechanically regulated, however the actual mechanical properties of the follicle microenvironment have remained unknown. Here we map and quantify the mechanical microenvironment in mouse ovaries using colloidal probe atomic force microscope (AFM) indentation, finding an overall mean Young’s Modulus 3.3 ± 2.5 kPa. Spatially, stiffness is low at the ovarian edge and centre, which are dominated by extra-follicular ECM, and highest in an intermediate zone dominated by large follicles. This suggests that large follicles should be considered as mechanically dominant structures in the ovary, in contrast to previous expectations. Our results provide a new, physiologically accurate framework for investigating how mechanics impacts follicle development and will underpin future tissue engineering of the ovary.

## Introduction

The ovary must precisely regulate the activation and development of follicles to release, in humans, exactly one oocyte (egg) per cycle. In contrast follicle dysregulation leads to disease and compromised fertility. In particular in polycystic ovary syndrome (PCOS), follicle development is impaired with consequent reduction in the rate of ovulation (1,2), while in the other direction uncontrolled and accelerated follicle development depletes follicle stocks leading to premature ovarian insufficiency (POI). In recent years it has been proposed that the mechanical properties of the ovarian environment are key to regulating follicle growth, in combination with previously identified factors of pituitary hormones, steroid hormones and growth factors (3–7). There is increasing evidence that the follicle responds to its mechanical environment, which either suppresses or enhances growth, with this mechanical environment controlled in turn by hormones or growth factors.

Evidence for mechanical regulation in follicle development includes on a fundamental cellular level the known role of the mechanosensitive Hippo (8,9) and Akt (10–13) signalling pathways. The most compelling evidence for mechanical control of follicle development, however, comes from experiments that have cultured follicles *in vitro* in biomaterial gels of controlled stiffness(14–19). These studies have shown that varying the gel stiffness can control follicle properties including growth, steroidogenesis and antrum formation, with a postulated switch from suppressive to a permissive environment for follicle development when the microenvironment stiffness drops below a threshold value in the range range 0.1 – 1 kPa (Young’s Modulus) (14,15,19). These experiments, combined with the knowledge that ECM content varies spatially through the ovary (20–23), have led to the hypothesis that follicle development is mechanically regulated by the different mechanical microenvironments of different ovarian regions (15,24–26). Since extra-follicular ECM density also varies dynamically across the menstrual cycle such ECM-driven mechanical control is a plausible means of dynamically regulating follicle development (27–30).

Despite these exciting concepts derived from in vitro culture, the actual mechanical properties of the follicle microenvironment, and how these vary spatially within the ovary remain unknown. Indeed the only substantial mechanical characterization of the ovary to date is a recent study that measures the bulk stiffness of the whole ovary using external indentation (31). Biomechanical control of follicle development in the actual ovary hence remains hypothetical. Here, we map the mechanical properties of the mouse ovarian interior with high spatial resolution, directly revealing the mechanical conditions that follicles will experience in the ovarian microenvironment. We use colloidal probe atomic force microscopy (AFM) indentation (32–36) to generate a spatial profile of stiffness (Young’s modulus) across a bisected ovary. We find a mean ovarian stiffness of 3.3 ± 2.5 kPa, with local fluctuations attributable to microstructural variation, but importantly also with substantial structural variation across the ovary. In contrast to prior expectations, the ovary is relatively soft at the edge and centre, regions that are dominated by extra-follicular ECM rich in collagen IV. However, in an intermediate zone between the centre and edge, where the tissue is dominated by large follicles, the ovary is much stiffer. This indicates that large follicles themselves are the dominant mechanical objects in this system, and focuses attention on direct follicle-follicle interactions as a likely key factor in follicle mechanoregulation.

## Results

We measured the local stiffness (characterized as Young’s modulus, *E*), using colloidal probe indentation with the AFM (for a full description see materials and methods). Briefly, a mouse ovary (25-29 days post-partum) was collected and embedded in an agar gel. The ovary and gel were then bisected with a sharp microtome blade to reveal the interior surface of the ovary (Fig 6). Using AFM, the cut surface of the ovary was probed at intervals from edge to edge across the ovary, using a spherical colloidal probe (10.8 μm diameter) mounted on the AFM cantilever. Colloidal probe indentation was selected to generate a reliable mechanical measurement since the Young’s modulus value will be averaged over an area of the same order of the sphere diameter, avoiding highly-localized effects. Equally, the ~ 10 μm probe size is much less than the ~ 2 mm size of the ovary itself, enabling a useful mapping of variation within the ovarian interior. At each site, a force-indentation curve was measured, and the Young’s modulus was extracted via a fit to the Hertz model (see Materials and Methods) (37). The fit was made to data at moderately high indentation, again to generate a reliable value for by averaging over a sufficiently large tissue volume, and to avoid any artefacts due to possible surface damage during ovary bisection.

Young’s modulus was mapped in a line-section across the ovary, illustrated in Figure 1D. The line-section was constructed by selecting locations spaced 50 – 100 μm apart across the ovarian interior surface and passing through its centre (insofar as this could be achieved by eye). At each location, a map of Young’s modulus was created 30 × 30 μm (6 × 6 grid of indentations with 5 μm spacing). This procedure reflected the construction of the AFM, which combined a 30 × 30 μm local piezo drive, and a micrometer-driven sample stage for larger spacing. To demonstrate reliability and technical reproducibility, the line-section was constructed to incorporate indentation at several locations on the stiffer agar gel surrounding the ovary, at both the start and the end of the section.

**Figure 1.**
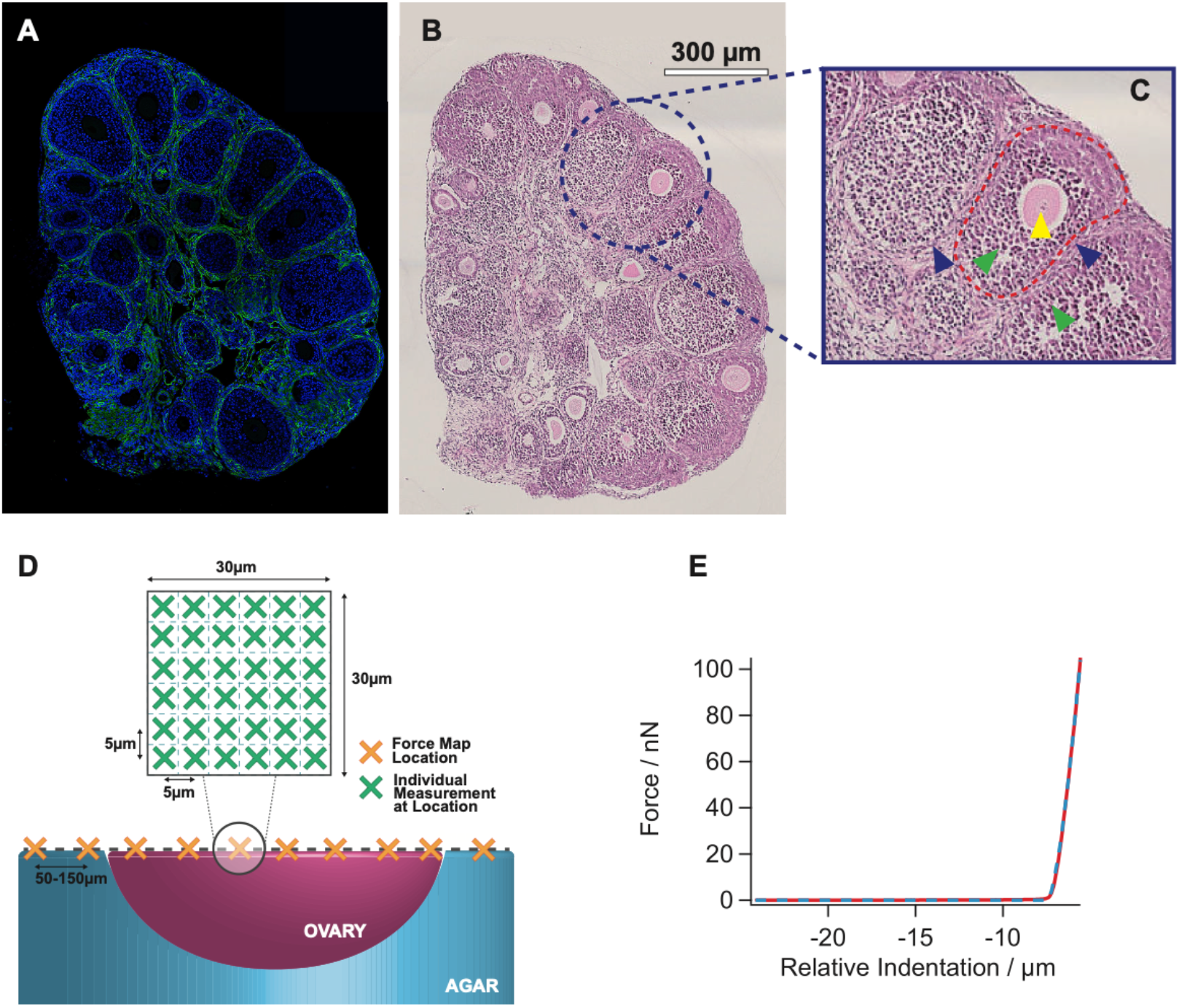
Measuring the mechanical properties of the ovarian microenvironment. (A,B) Representative histology images showing the exposed surface of a bisected ovary, on which the mechanical indentation measurements were performed. Staining is (A) Immunofluorescence showing collagen IV as a marker for ECM (green) and DAPI staining of cell nuclei (blue). (C) Closeup showing a follicle (outer border given by red dotted line), indicating the oocyte (indicated by yellow triangle), granulosa cells (green triangle), theca cell layer (blue triangle). D)Schematic of indentation line-scan: a series of locations across the ovary interior surface (X) with spacing 50 – 100 μm are selected. At each location, a 30 × 30 μm map is created by indenting at a 6 × 6 grid of sites (solid dots) with 5μm spacing. (E) A representative curve of force, F, vs indentation δ (the values on the x-axis are relative since the surface location δ_0_ is unknown until after fitting) curve, showing a fit to the Hertz model using data at moderate indentation as per the methods section.

The stiffness of agar was consistently of order 15 kPa, and most importantly showed no appreciable alteration between the start and end of the line-sections (Figure 4A). To confirm repeatability and that indentation does not significantly damage the tissue surface, we constructed multiple maps at the same location, showing no significant difference in the values recorded in first and second indentation (p > 0.05, Supplementary Figure 1).

### The ovary is a fairly soft tissue but with very broad micromechanical variation

We first considered the overall distribution of the Young’s modulus values measured (n=4 ovaries, Fig. 2, Left Panel). It can be seen that a substantial range of values (c. 0.5 kPa to 10 kPa.) is measured, summarized by a mean value of 3.3 ± 2.5 kPa or median 2.6 *±* 2.8 kPa (Fig 2, Right Panel) (where *±* signifies standard deviation and interquartile range respectively). Interestingly, the distribution is heavily skewed towards low stiffnesses, with a large number of values below 1 kPa. Comparing the mean and standard deviation values we have measured for the ovarian stiffness with those measured for other tissues (32,33,38–41), (Figure 3), we see that the ovary emerges as an overall fairly soft tissue, comparable with the kidney or fat, but not as soft as e.g. the brain. These overall values are in good accordance with recent measurements of whole-ovary stiffness using exterior indentation, which quoted c. 2 kPa for ‘reproductively young’ mouse ovaries that are roughly comparable with those measured here. However looking in more detail reveals a more complex picture (31). The ovarian microenvironment is seen to range from very low stiffnesses usually associated with the brain as the softest tissue up to values that approach those associated with stiff tissues such as muscle. Hence it becomes clear that mechanical variations within the ovarian microenvironment may be as significant as variations between the ovary and other tissues. As a further contextualization, we compare our measured values for the ovarian stiffness with the permissive stiffnesses for follicle activation in controlled-stiffness gels discussed above (14–19,42,43). In this work, examples of Young’s modulus values considered permissive for follicle development are in the range 100 – 800 Pa (Supplementary Table 1) (14,15,19), however the upper limit is not necessarily precisely defined. Given that the ovary does indeed exhibit a substantial number of sites where stiffness is at sub-kPa values, it is possible that the requirements for ‘permissiveness’ could be fulfilled.

**Figure 2:**
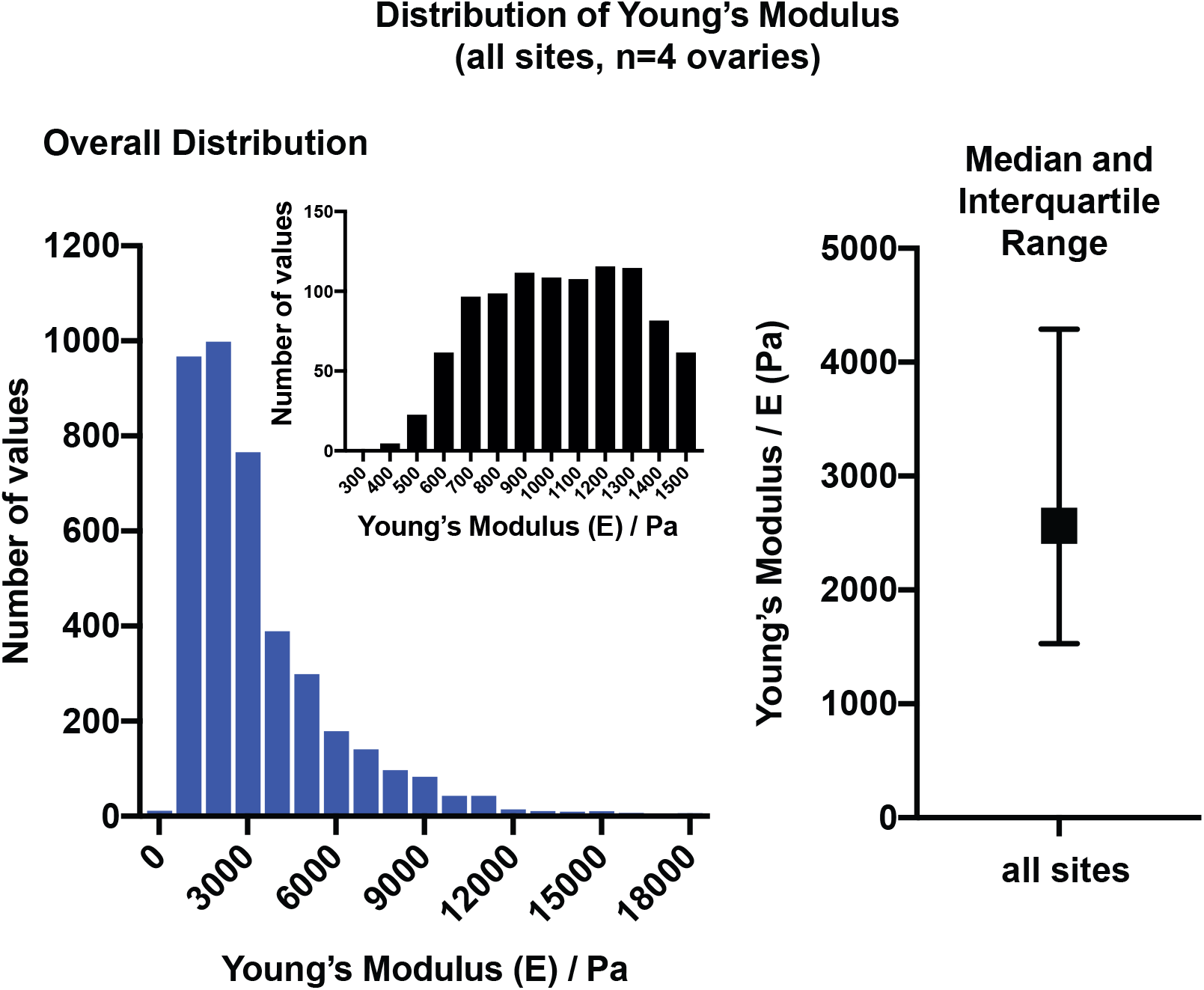
(Left panel) A histogram representing the overall distribution of Young’s moduli (E)/ Pa across all probed ovaries (within culture media, n=4 ovaries). X-axis values correspond to upper limit of bin, e.g. 3000 Pa implies values between 2000 and 3000 Pa. Each value within the histogram represents one force curved produced. Inset shows the low-stiffness part of the histogram at higher resolution (smaller bin size). (Right panel) Median and interquartile range of the data across all samples.

**Figure 3:**
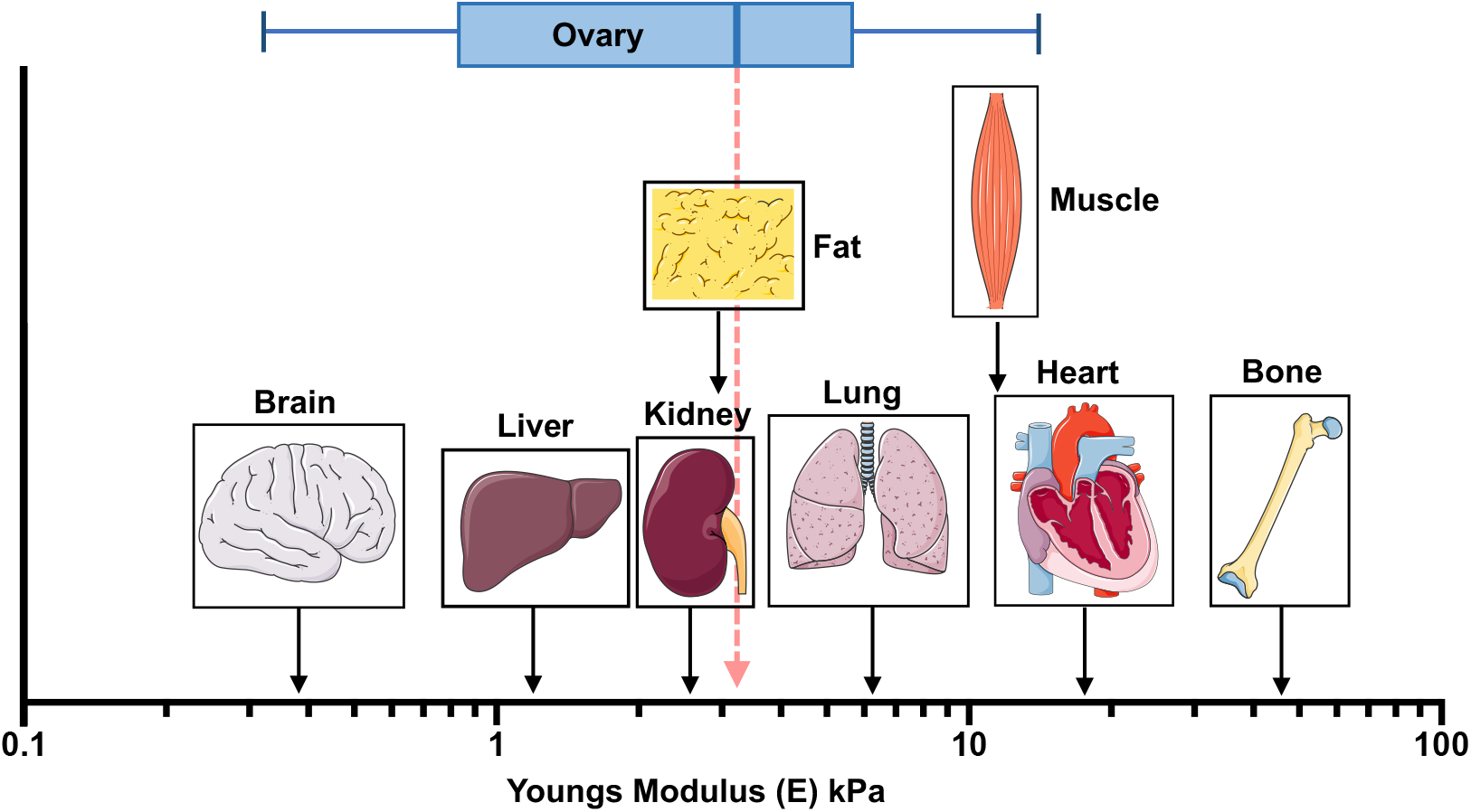
The measured stiffness of the ovary placed in the context of other mouse tissues (data for other tissues reproduced and figure adapted from (66)).

### The ovary is softest at the centre and edge, with an intermediate zone of maximal stiffness

Line profiles of stiffness across the ovaries were plotted (Figure 4). The raw data for a representative example (Figure 4A) shows how the softer ovary sits within the stiffer agar gel used as a matrix. We then extract the ovary data alone (representative example Figure B), displaying the mean and standard deviation of Young’s modulus values found within the local stiffness map at a specific location (X in the schematic Figure 1D). The experiment was repeated across a total of seven ovaries from different animals (Figure 4B, 4C). A characteristic double-peak structure was identified and we now discuss this.

**Figure 4.**
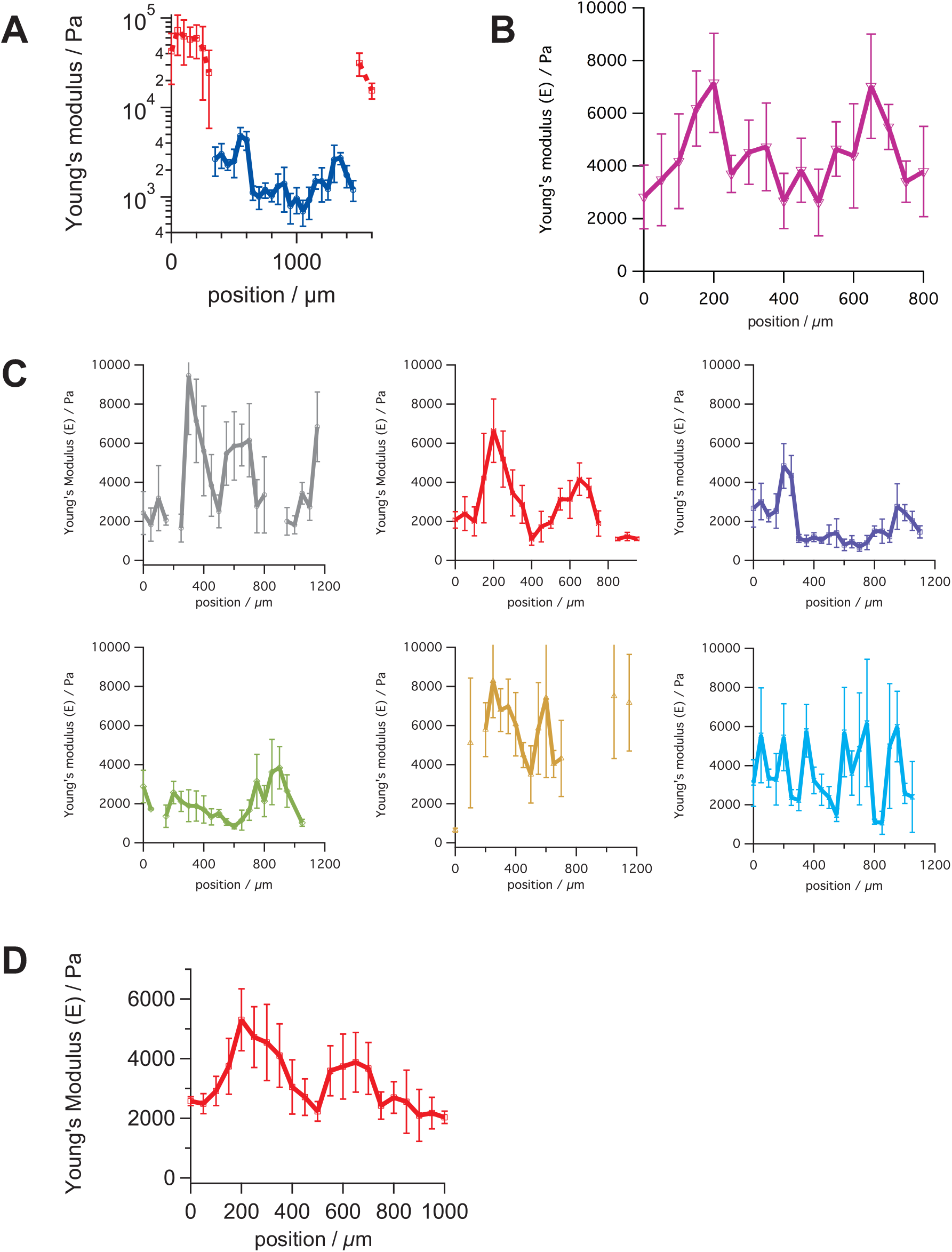
Line-scan stiffness (Young’s Modulus, E) profiles across ovaries. (A) Representative line-scan, showing the softer ovary (blue) embedded within the stiffer agar gel (red). Markers indicate mean and bars standard deviation of the stiffnesses measured within the local stiffness map at each location. (B) A representative example of an ovary profile. (C) Linescans across other ovaries, making a total of 7 ovaries scanned, of which 6 show the characteristic double-peak structure discussed in the text. (D) Averaged line-stiffness profile (mean of profiles that match double peak characteristic, bars indicate standard deviation).

The edges of the ovary are softer, with a Young’s Modulus that varies between ovaries but is in the region of 2 – 3 kPa. Moving towards the centre of the ovary, the stiffness rises to a peak which again varies but is significantly greater, with an average peak value of ~ 7 kPa (Figure 4B). This peak stiffness does not occur at the centre of the ovary, rather twin peaks are observed on either side of the centre, which itself is soft, similar in stiffness to the edge region. This double-peak structure of the stiffness linescan profile is observed in 6 out of 7 ovaries (Figure 4C). Look in particular at the averaged line-scan (Figure 4D, created by taking the mean of the ovary profiles). Hence, we can give a well-defined description of the microenvironment that will be experienced by follicles throughout the ovary.

It is interesting to note that the observed spatial variation of stiffness within the ovary had not been previously predicted, emphasizing the importance of directly mapping tissue stiffness using approaches such as AFM.

### Higher-stiffness regions in the ovary are associated with larger follicles, and not with inter-follicular ECM

Stiffness in tissues is often associated with ECM components, notably collagens. A common model is that collagens are primarily responsible for the stiffness of tissues with a high collagen concentration typically thought to correspond to a stiff microenvironment and vice versa (44). Indeed this concept has been applied to the ovary by several authors who have postulated that the high concentration of collagen often seen at the edge [cortex] of the ovary should make it the stiffest region of the ovarian microenvironment (24,45). However we have seen clearly from our AFM measurements that directly characterize mechanics that the edge of the ovary is not the stiffest region. Hence we now examine quantitatively whether the stiffer regions of the ovary indeed correspond to the highest collagen concentration, or whether other mechanisms are at work.

Following AFM analysis, the probed bisected ovary was fixed in formalin, paraffin embedded and sectioned. Collagen IV, which is highly expressed in the basal lamina and theca layer (46) was localized in the probed section (Figure 1A, representative image). To generate a clear idea of the quantity of collagen IV in the different parts of the ovary, used as a marker for the inter-follicular space, we determined the average amount of collagen IV per unit area as a function of distance from the ovarian center. Mathematically, we calculated the two-dimensional radial distribution function *g(r)* of the collagen IV immunofluorescence intensity, defined as the intensity per unit area in an annulus [ring] whose outer edge is distance *r* from the ovary center (Figure 5A). The results are consistent across ovaries (Figures 5B,C). There is a region of higher collagen IV high at lower values of *r*, near the ovary center. Another region of higher collagen IV density is seen near the ovary edge. In between, there is a region of lower collagen IV density.

**Figure 5.**
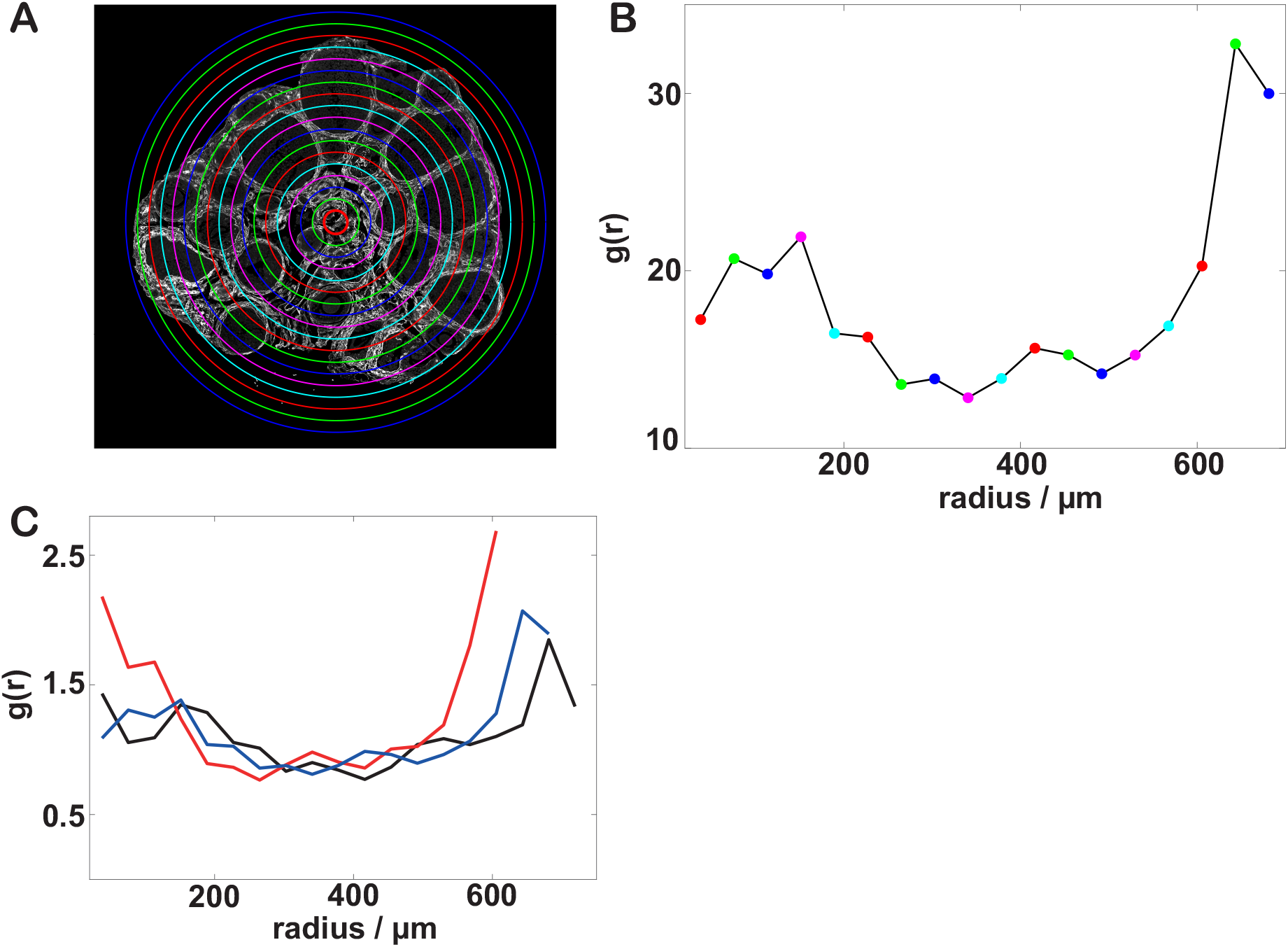
The distribution of collagen IV within the ovary. (A) Annuli used to calculate the radial distribution function for a representative ovary, imposed on the collagen IV immunofluorescence histology image. (B) The radial distribution function g(r) of collagen IV immunofluorescence intensity, calculated using (A). (C) g(r) for 3 representative ovaries.

It is hence immediately clear that the regions of high collagen IV density do not correspond to the regions of high stiffness identified above. Rather there is an inverse relationship. The regions of higher collagen IV density at the center and edge are also regions of lower Young’s modulus, while the region of lower collagen IV density in between the two corresponds to a peak in Young’s modulus.

Looking in detail at the immunofluorescence images, we see why the overall collagen IV density is low in the high-stiffness region: this area is dominated by larger follicles, with the follicular interior consisting of granulosa cells which express little or no collagen IV, the oocyte and, in some larger follicles, a fluid-filled antrum (a cavity formed in the granulosa cell layer). For the specific case of the ovary, therefore, it seems that the mechanical properties of these large follicles, rather than of the extra-follicular ECM, is the dominant factor in determining which areas of the ovary exhibit high vs low stiffness. Note that these results apply to reproductively young ovaries as measured here: it is possible that age-induced fibrosis may develop a greater role for inter-follicle collagens as the ovary ages (31).

While the dominance of follicles as drivers of high stiffness is clear from our results, the microscopic origin of this stiffness may be attributable to a range of different components that make up the follicle. The oocyte itself has been measured to have a high stiffness (47), although this may depend strongly on developmental stage. Equally, the follicle itself contains some collagen I (20). F-actin is likely to participate, given the observation of highly-defined rings of F-actin rich cells surrounding the oocyte and in the theca layer (4). Finally, follicles as growing tissues, will intrinsically build up an internal pressure (48) rendering them prestressed and difficult to compress further.

### High resolution measurements show mechanical microstructuring of the ovary on a local scale

Having considered how the mechanical environment varies on a large scale across the ovary, we now consider local variations. Recall that, at each location within the ovary a 30 x 30 μm grid of measurements was mapped with up to 36 indentations, we now proceed to consider the variation within the maps themselves (rather than simply the map average plotted in Figure 4). Taking a simple approach, we can see comparing different locations that the standard deviation of Young’s Modulus as a fraction of the mean for each location remains roughly constant at roughly one third across many locations and several ovaries (Figure 6B). That is, the stiffest locations showed the most variation. Plotting selected local stiffness maps (Figure 6A): it is clear that the locations of greatest stiffness (the peaks on the trans-ovary profiles (Figure 4)), exhibit very substantial variation. For example in the peak-located heat map 6Aii we see a full stiffness range of almost 8 kPa. In comparison to the gap between mean stiffnesses at peak and trough, which is roughly 4-5 kPa (see above), the local variation is hence smaller but still very significant. Looking at the local stiffness maps themselves, we can see that higher and lower Young’s Modulus values are not randomly distributed within the maps but rather there are clear spatial correlations, i.e. there are clear regions of higher and localized stiffness within the maps. This indicates the presence of mechanical microstructure, i.e. regions of high and low stiffness can be discerned on the ~ 10 μm lengthscale.

**Figure 6:**
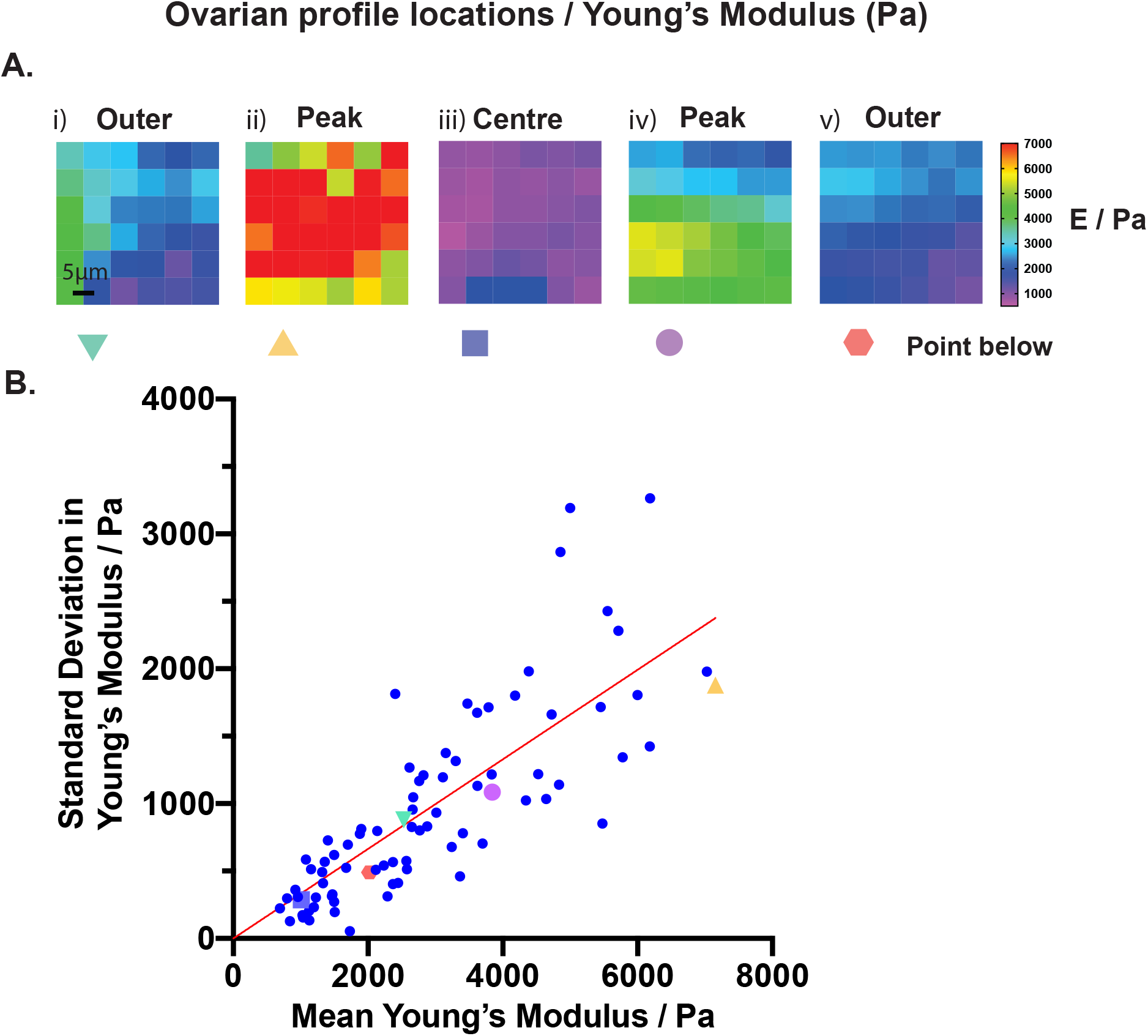
Mechanical microstructuring. A. Demonstrates a range of heatmaps over various spatial locations along the ovary transept. Each square represents the modulus taken at that site. Values over 7 kPa were truncated for scaling. B. Represents the mean Young’s modulus of each site against the standard deviation of that site.

**Figure 7:**
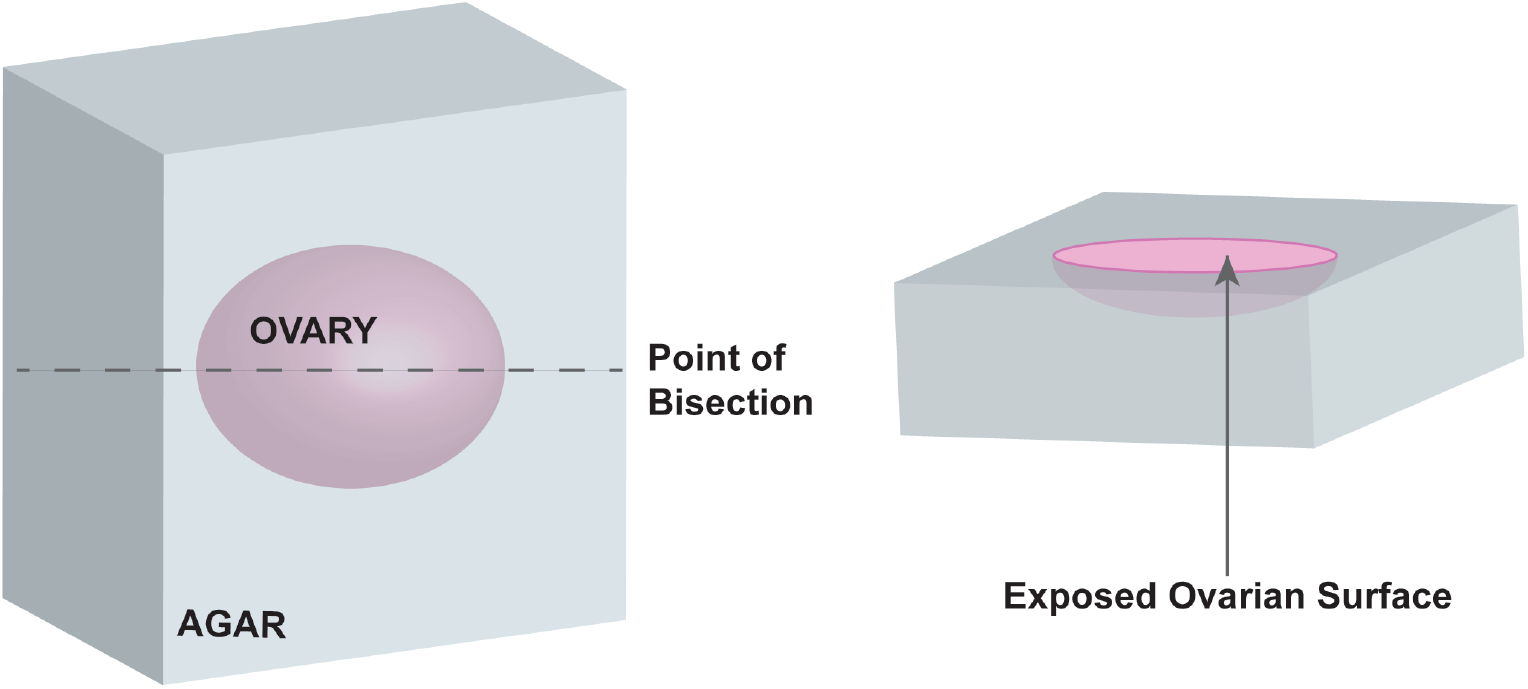
Representation of the technique required to prepare the sample before measurement. The ovary is immersed within a 2% agar solution and bisected, exposing its interior for analysis, prior to being placed within the AFM.

In contrast to the large-scale variations in stiffness that we observe across the ovary (Figure 2), this local microstructure is clearly not due to the presence or absence of larger follicles, since these are typically ~ 100 μm or more across. Rather these variations must represent microstructural variations within the ECM, between ECM or cellular components in the follicle.

Considering the history of tissue engineering and biophysics research both in the ovary and other tissues, a valuable simplification has often been to culture cells and tissue constructs within bioengineered gels whose mechanical properties are presumed homogeneous, with each gel characterized by a single Young’s modulus. While unsurprising, it is intriguing to consider that in real tissue, microstructural variations in mechanics may be very substantial. We do not know how tissues respond to mechanically microstructured materials and how this may differ from their responses in the homogeneous case.

## Conclusion

We have measured for the first time, and mapped spatially, the mechanical properties of the mouse ovary interior. The Young’s Modulus ranges between approximately 0.5 – 10 kPa, with substantial spatial variation on a microstructural level, in addition to larger-scale differences between the different regions of the ovary. This complex structuring emphasizes that tissue components such as cells and follicles experience a complex mechanical microenvironment that cannot be reduced to a single number such as a tissue-scale Young’s modulus. Analyzing large-scale spatial variations across the ovary shows lower stiffness regions at the ovarian edge and centre, with regions of peak stiffness in between. The areas of low stiffness coincide with a high density of collagen IV, contradicting previous assumptions that the ovarian microenvironment is mechanically dominated by extra-follicular ECM. Rather the higher stiffness regions are dominated by large developmentally advanced follicles, suggesting that these structures should be considered mechanically dominant. The mechanical importance of these follicles stands in contrast to the widespread reductive assumption that stiffer regions can be identified by considering only one component such as collagens. The mechanical characterization of the ovary will be pivotal in understanding how mechanical effects impact follicle development and the development of diseases including PCOS and POI. Equally, a knowledge of the real mechanical properties of the ovarian microenvironment will guide the design of ex-vivo tissue-engineered mimic ovaries.

## Materials and Methods

### Ovary Collection

Whole ovaries were collected from female C57BL/6 mice pups (Charles River) euthanased at days 25-29 post-partum. Mice were housed in accordance with the Animals (Scientific Procedures) Act of 1986 and associated Codes of Practice.

#### Atomic Force Microscope (AFM) indentation measurements

The ovaries were removed and cleaned of extraneous tissue (uterus, oviduct, fat) using insulin needles over a heated microscope stage (37°C) in L-15 isolation media (Life Technologies, Paisley, United Kingdom) supplemented with 1% (weight/volume) bovine serum albumin (Sigma). A 2% agarose (Sigma) solution was created and allowed to cool before being placed into a glass cavity block (40mm x 40mm). The ovary was immersed and positioned within the agarose and allowed to solidify completely. The agarose platform was then dissected through the centre of the ovary using a microtome blade (MX-35, Thermo-Fisher) and glued (Loctite) to the base of the heated bio-cell.

An atomic force microscope (Asylum MFP-3D, Oxford Instruments) mounted on an incorporated inverted light microscope was used to measure material properties by indentation across a transverse section of the ovary. The colloidal probe used for all indentations had a spring constant, *k*, of between 0.02 – 0.77 N/m with a 10.8μm polystyrene sphere attached to the tip of the probe (sQUBE, Bickenbach, Germany). A fluid cell was used to maintain samples within culture medium or water at a constant 37°C. Before each experiment the inverse optical lever sensitivity (Invols) in water was calibrated by collecting a force curve on a hard surface (glass). The spring constant was determined on the thermal noise spectrum using the instrument manufacturer’s software.

Samples were equilibrated on the heated AFM stage at 37°C, under L-15 cell culture medium or deionized water (MilliQ), before data collection occurred. Using the microscope viewing display, the probe was lowered over a region of agarose close to but outside the tissue itself. Using the built in micrometer-driven sample stage, indentation-based mapping of mechanical properties was then carried out at a series of locations, 50 – 100 μm apart along a diameter across the cut exposed ovary surface. At each of these locations, indentation maps were created, consisting of 36 measurements arranged in a 30 x 30μm square grid with 5μm spacing between measurement sites, with movement within the grid carried out using the AFM piezoelectric drive. To determine repeatability, 3 indentation maps were identically measured at each location.

### AFM data analysis and parameter extraction

The measured indentation curves were analysed to extract the value of Young’s modulus. Approaching curves only were analysed, and a custom-built software programme was created in Igor Pro (Wavemetrics Inc.) to treat the data as follows. Initially, each force-extension curve was accepted or rejected based on requiring a minimum of 10 μm of overall travel, to exclude curves where the out-of-contact baseline was insufficient. A second criterion of requiring a minimum of 300nM of cantilever deflection was added as the project progressed. The user was also given the chance to manually reject the few curves where, for example, the out-of-contact baseline was curved due to varying drift. To remove the effect of typically small linear drifts, a linear baseline fitted to the out-of-contact region was first subtracted before the force-extension curves were converted into force-deflection curves using the measured value of the cantilever spring constant. The Young’s modulus was determined by a Hertz model fit, made in the region of moderately deep indentation (Figure 1E). This was done to exclude the region of initial contact where the apparent stiffness might be affected by surface damage due to the microtome blade. In addition, measuring at a reasonably deep indentation means that the measured Young’s modulus is in effect integrated over a larger volume of the sample, implying greater reproducibility. The equation used for fitting was

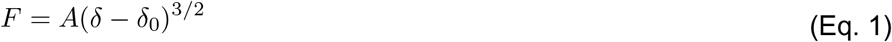

where *F* is the force, (*δ − δ*_0_) the indentation with *δ*_0_ being the surface position, and A a prefactor given by (37).

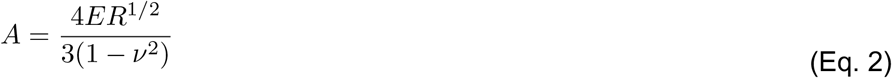

The free parameters *A* and *δ*_0_ were determined by least squares fitting (Igor Pro intrinsic function) and then *E* was determined using the known value of *R* and an assumed Poisson’s ratio of 0.45, representative of the values quoted for soft tissues (48,49).

The repeatability of the measurements was confirmed by comparing the multiple maps performed at each site. No significant difference was seen between the statistical distribution of Young’s Moduli first and second mappings at a given location (Supplementary Figure 1A) and comparing the mappings shows an essentially unchanged picture also when taking spatial location into account (Supplementary Figure 1B). This confirms reproducibility and shows that indentation does not appreciably damage the tissue surface. Once this had been confirmed, first mappings (ie. The first indentation at each site) were used throughout except in some cases where the number of rejections was too high to deliver sufficient good force curves from separate sites in each map, in which case values from first and subsequent mappings were combined to satisfy this criterion.

### Histology

Upon completion of experiments the tissues were fixed in Formalin (10% Neutral Buffered) (HT5014, Sigma-Aldrich) overnight. Using a microscope stage (Nikon), bisected ovaries were arranged so that the probed surface faced downwards and set in 2% agar (Sigma). Samples were processed through graded alcohol (70% 1h), 90% (1h), 100% (3 × 1h)) and then placed in Histoclear (Cat. No HS-200, National Diagnostics) overnight, followed by a further 1h the following morning. Samples were then embedded in paraffin wax for 2 hours at 55°C, at the correct orientation so that the probed ovarian surface could be analysed.

Formalin fixed (HT5014, Sigma-Aldrich), paraffin embedded ovaries were serially sectioned (5μm) parallel with the cut surface using a Leica RM2135 microtome (Leica Microsystems UK Ltd. Milton Keynes UK) and mounted onto slides (SuperFrost Plus, VWR International Ltd.)

### Haematoxylin and Eosin Staining

Samples were sectioned and stained using haematoxylin and eosin (H&E), following the standard protocol. Briefly, the tissue sections were dewaxed using xylene (Scientific Laboratory Supplies) and hydrated by passing through decreasing concentration of alcohol (100%, 90%, 70%). Following staining using Gills haematoxylin () for 3 minutes and washed under running tap water. The sections were then differentiated by submersion in 1% acid alcohol (1% HCL and 70% alcohol) and again washed in water. 1% Eosin was applied for 5 seconds followed by dehydrated in increasing concentration of alcohol and cleared in xylene before mounting with mounting media (Eukitt, Scientific Laboratory Supplies).

### Immunofluorescence imaging

Slides with central sections of ovary were de-waxed in Histoclear (National Diagnostics) for 5 mins x 2, then re-hydrated in 100%, 95% and 70% ethanol. They were finally washed in deionised water for 2 x 5 mins. Slides were boiled for 20 mins in a citrate buffer for antigen retrieval and left to cool for 30 mins. Slides were consequently washed in phosphate buffered saline (PBS) for 3 x 10 mins. To reduce non-specific binding, slides were blocked with 5% normal goat serum (heat-inactivated) or 10% normal donkey serum in PBS, supplemented with 4% BSA () for 30 mins at RT. Primary antibodies were diluted in either 5% or 1% serum/BSA in PBS and applied to the sections overnight at 4°C.

The following day, slides were washed in PBS 3 x 10 mins. A secondary antibody conjugated to Alexa-Fluor™ 488 (Abcam, A150077) was added to all slides, for 1 h at RT, in the dark. After, slides were rinsed in PBS for 3 x 5 mins and stained with 1 %g/mL 4,6-diamino-2-phenylindole (DAPI - Molecular Probes) for 3 mins in the dark at RT. Slides were mounted in ‘Prolong Gold’ Antifade P36931 (Molecular Probes) containing DAPI and left at RT for 24 h before imaging. The slides were then imaged using a Leica inverted SP5 confocal laser-scanning microscope (Leica Microsystems, Wetzlar, Germany). Individual images were taken across the entire sample to generate a full ovarian cross-sectional image. In the ovaries selected for quantitative analysis, laser and detector settings were held constant across all samples and positions, and these ovaries were also stained simultaneously.

### Image analysis

Individual images were stitched together in two distinct ways. For pure visualization a manually-based stitching software DoubleTake (Echo One, Denmark) was used across all channels. For image quantification, scans were stitched together in FIJI using the function *pairwise stitching*. These images were segmented in MATLAB (The MathWorks, Natick, MA) using a graph cut segmentation, with morphological closing subsequently applied to smooth edges (MATLAB function *imclose*, disc radius 12 pixels). The centre of each ovarian section was calculated from the segmentation.

To quantify the distribution of collagen IV within the ovary, a radial distribution function was calculated, measuring the average intensity of collagen fluorescence in annular regions of width Δ*r* centred on each ovarian section. Specifically, for *r_i_*=*i* Δ*r*

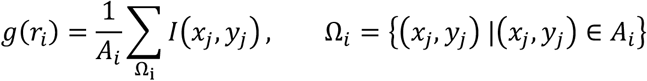

Where *I(x,y)* is the intensity and *A_i_* is the area of the annulus *r*_*i*_ − Δ*r* < *r* < *r*_*i*_ lying within the ovarian section thus adjusting for edge effects. Of course, where the annulus lies entirely within the tissue section 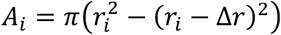. The annulus width Δ*r*=*37.8μ*m throughout.

## Acknowledgements

We acknowledge funding from the EPSRC (Doctoral Training Programme scholarship to TH). Additional consumable funds were kindly provided by the Genesis Research Trust (GRT).

## SUPPLEMENTARY INFORMATION

**Supplementary Figure 1:**
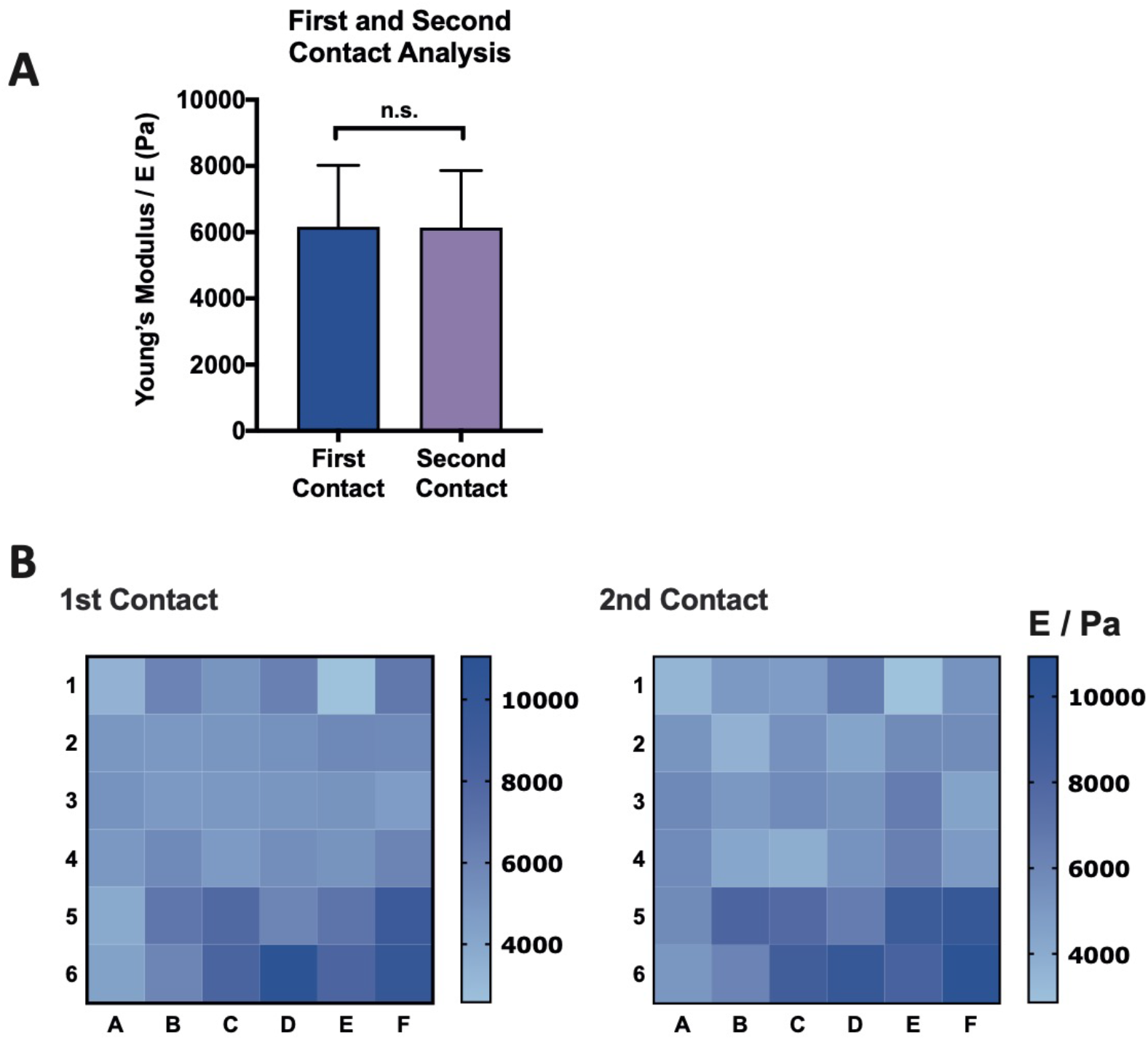
First and second compressions of ovaries show no statistically significant difference. A) Graph demonstrating a representation of the difference between first contact and second contact at the same site, there is no significant difference between the two groups demonstrating no change in the results taken from the first or second contact (P>0.05). Normality was assessed using a D’Agostino and Pearson test. A paired t test was performed. Graph shows mean and SD B) Heatmaps that show the Young’s Modulus at a single position at the first and second contact.

### Literature analysis of permissive versus suppressive controlled-stiffness gels for follicle development

**Supplementary Table 1:**
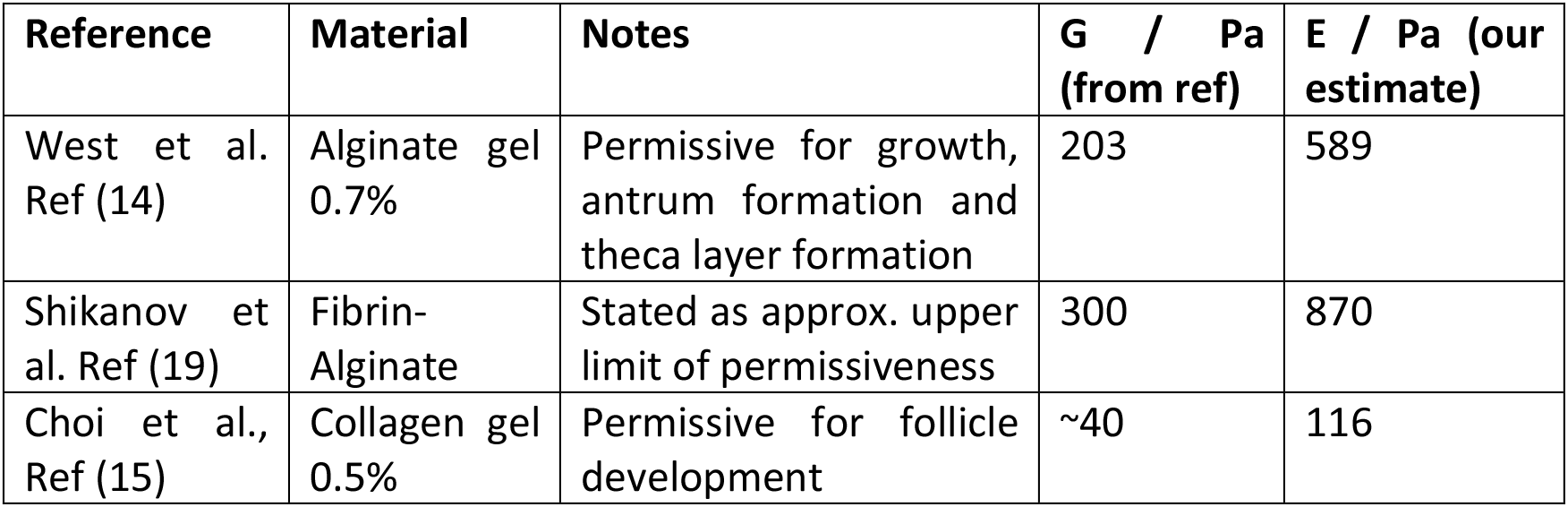
Literature analysis showing the stiffness of hydrogels defined as permissive for follicle development by various authors. Values quoted by these authors are shear modulus G (or alternatively storage modulus G’), converted to Young’s Modulus using E = 2G(1 + ν) (50) taking the Poisson’s ratio, v, as ~0.45 (49).

| Reference | Material | Notes | G / Pa (from ref) | E / Pa (our estimate) |
| --- | --- | --- | --- | --- |
| West et al. Ref (14) | Alginate gel 0.7% | Permissive for growth, antrum formation and theca layer formation | 203 | 589 |
| Shikanov et al. Ref (19) | Fibrin-Alginate | Stated as approx. upper limit of permissiveness | 300 | 870 |
| Choi et al., Ref (15) | Collagen gel 0.5% | Permissive for follicle development | ~40 | 116 |

**Supplementary Table 2:**
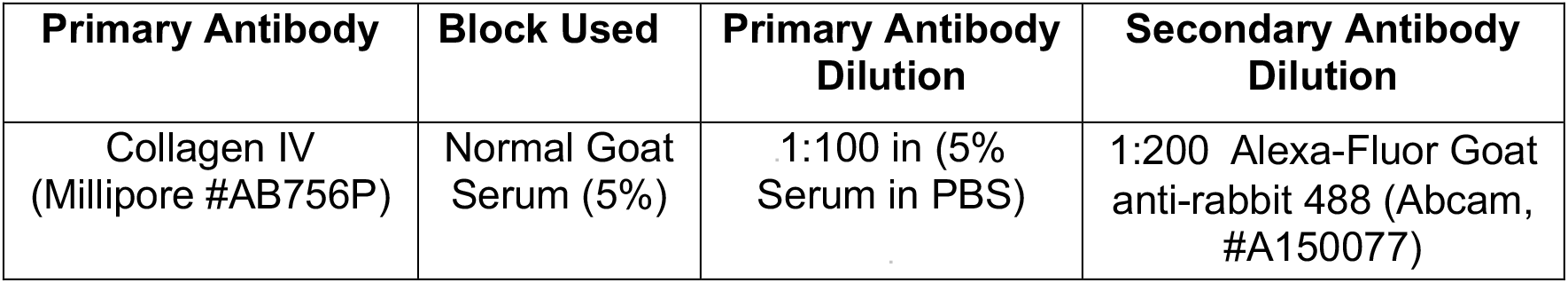
Antibodies used for Immunofluorescence.

| Primary Antibody | Block Used | Primary Antibody Dilution | Secondary Antibody Dilution |
| --- | --- | --- | --- |
| Collagen IV (Millipore #AB756P) | Normal Goat Serum (5%) | 1:100 in (5% Serum in PBS) | 1:200 Alexa-Fluor Goat anti-rabbit 488 (Abcam, #A150077) |

